# A robust method for generating, quantifying and testing large amounts of *Escherichia coli* persisters

**DOI:** 10.1101/2020.07.06.181891

**Authors:** Silke R. Vedelaar, Jakub L. Radzikowski, Matthias Heinemann

## Abstract

Bacteria can exhibit phenotypes, which makes them tolerant against antibiotics. However, often only a few cells of a bacterial population show such so-called persister phenotype, which makes it difficult to study this health-threatening phenotype. We recently found that certain abrupt nutrient-shifts generate *E. coli* populations that consist of almost only antibiotic tolerant persister cells. Such nearly homogeneous persister cell populations enable assessment with population-averaging experimental methods, such as high-throughput methods. In this paper, we provide a detailed protocol of how to generate such large fraction of tolerant cells using the nutrient-switch approach. Furthermore, we describe how to determine the fraction of cells that enter the tolerant state upon a sudden nutrient shift and describe a new way to assess antibiotic tolerance with flow cytometry. We envision that these methods facilitate research into the important and exciting phenotype of bacterial cells.

## 1. Introduction

Bacterial persistence is defined as the occurrence of cells within a population that are tolerant against antibiotics without carrying a genetic resistance (1). Such antibiotic tolerant cells were suggested to be responsible for recurrent infections (2). Tolerant cells can be formed stochastically in exponentially growing cultures (3). Activation of toxin and anti-toxin (TAS) modules (2) and deletion of certain metabolic genes (4) has also shown to increase the number of cells entering this phenotype. Certain environmental perturbations can also induce the fraction of tolerant cells in a population, such as the entry into stationary phase (5) or certain sudden nutrient shifts, where almost all the cells in a population can enter the tolerant state (6, 7).

The fact that certain nutrient shifts, in *E. coli*, for instance, the one from glucose to fumarate (8), can force almost all the cells in a population into the tolerant state offers new research opportunities to investigate the molecular basis of tolerance and persistence. While the investigation of the few stochastically occurring persisters in exponentially growing cultures and the very heterogeneous stationary phase cultures require single-cell analyses or cell-sorting approaches, a population that consists of almost only tolerant cells allows the use of population-level high-throughput analyses. For instance, exploiting sudden nutrient shifts to generate almost homogeneous populations of tolerant cells, it was found that tolerant cells have increased ppGpp levels, are metabolically active with a metabolism geared towards energy generation and catabolism, and exhibit a proteome characterized by σ^S^□mediated stress response (8).

Despite the now enabled use of omics techniques for the study of bacterial persistence, experiments to assess how many cells are tolerant at which level of antibiotic exposure are still necessary. Here, in most cases, classical plating assays, using different dilutions, to determine the colony-forming units are performed (e.g. (5, 6)). However, this technique is very laborious and it has a high experiment-to-experiment variability requiring many plates to obtain statistically sound results (9). Moreover, with plating fewer cells are recovered from their dormancy compared to recovery in liquid media (10). To this end, we recently introduced an alternative method to assess antibiotic tolerance with flow cytometry (8). This method can be applied to populations solely consisting of dormant cells or to heterogeneous populations with dormant and growing cells. In this method, single cell’s regrowth is assessed via temporal dilution of a fluorescent signal, which can originate from a stained membrane (11) or GFP expression (12).

Here, we provide a detailed nutrient-shift-based protocol to generate large fractions of tolerant cells, which enables the study of tolerant cells on the population level. Further, we illustrate a method to fluorescently stain the membrane of cells, which, together with flow cytometry and a Matlab script, allows us to determine the fraction of cells that enter the tolerant state upon a sudden nutrient shift. Finally, we describe how the membrane staining procedure together with flow cytometry can be used to assess tolerance.

## 2. Materials

Prepare all media using demi water.

### 2.1. The generation of large fractions of tolerant cells

1. LB agar plates: prepare LB medium by adding tryptone (to a final concentration of 1% w/v), yeast extract (0.5% w/v) and sodium chloride (1% w/v) to a bottle with demi water. Add agar to a final concentration of 1.5% (w/v) and mix well. Autoclave for 15 minutes at 121°C and let cool down to ~55°C. Add antibiotics, if needed, and pour ~20 mL LB-agar per 10 cm petri dish (*See* **Note 1**). Store plates sealed with parafilm at 4°C with the agar side up. LB-plates containing antibiotics can be stored for up to one month.
2. M9 minimal medium: Prepare a base salt solution (211 mM Na_2_HPO_4_, 110 mM KH_2_PO_4_, 42.8 mM NaCl, 56.7 mM (NH_4_)_2_SO_4_), a solution with trace elements (0.63 mM ZnSO_4_, 0.7 mM CuCl_2_, 0.71 mM MnSO_4_, 0.76 mM CoCl_2_), a solution with 0.1M CaCl_2_, one with 1 M MgSO_4_, one with 1.4 mM thiamine-HCl, and one with 0.1 M FeCl_3_. Autoclave the solutions with base salts, CaCl_2_, MgSO_4_, and trace elements, and store them at room temperature. Sterile filter the thiamine and FeCl_3_ solutions using a 0.22 μM PES filter (*See* **Note 2**) and store them at 4°C. Stock solutions are sterilized so they can be stored for up to half a year. For 1 L M9 minimal medium, add 200 mL base salts, 700 mL water, 10 mL trace elements, 1 mL CaCl_2_ solution, 1 mL MgSO_4_ solution, 0.6 mL FeCl3 solution and 2 mL thiamine solution. Fill up to 1 L with water and filter sterilize the resulting solution using a PES bottle top filter into an autoclaved 1 L bottle. Store at 4°C for up to one month. The required carbon source is added fresh before the medium is used for cultivations.
3. 250 g/L glucose stock solution: For 100 mL, weigh 27.5 g D-glucose monohydrate and dissolve in 90 mL water. Adjust pH to 7 using NaOH and fill up to 100 mL with water. Filter sterilize the resulting solution using a PES bottle top filter into an autoclaved 100 mL bottle. Store at room temperature for up to one month.
4. 100 g/L fumarate stock solution: For 100 mL, weigh 14 g sodium fumarate and dissolve in 90 mL water. Adjust pH to 7 using HCl and fill up to 100 mL with water. Filter sterilize the resulting solution using a PES bottle top filter into an autoclaved 100 mL bottle. Store at room temperature for up to one month.
5. M9 minimal medium supplemented with glucose to a final concentration of 5 g/L for immediate use for cultivation: Add 1 mL glucose stock solution (250 g/L) to 49 mL minimal medium in a 500 mL Erlenmeyer flask. Preheat and pre-aerate at 37°C with shaking at 300 rpm before inoculation.
6. M9 minimal medium supplemented with fumarate to a final concentration on 2 g/L for immediate use for cultivation: Add 1 mL fumarate stock solution (100 g/L) to 49 mL minimal medium in a 500 mL Erlenmeyer flask. Preheat and pre-aerate at 37°C with shaking at 300 rpm before inoculation.

### 2.2. Identification and quantification of tolerant cells using membrane staining and flow cytometry

1. M9 minimal medium without carbon source. Ice-cold.
2. M9 minimal medium with 1% (w/v) bovine serum albumin: dissolve 0.5 g bovine serum albumin in 50 mL M9 minimal medium and filter sterilize the resulting solution using a PES bottle top filter into an autoclaved 100 mL bottle. Store at 4°C.
3. Dye to stain cell membrane: PKH67 dye (, Sigma), keep at 4°C until use. Then dilute 50x in Diluent C, at room temperature.
4. Diluent C (Sigma, included in the PKH67 kit). Store at 4°C, let warm up to room temperature before the start of the staining.
5. M9 minimal medium without carbon source, at room temperature.
6. M9 minimal medium with fumarate added to a final concentration of 2 g/L, preheated to 37°C, and pre-aerated.

### 2.3. Assessment of antibiotic tolerance with flow cytometry

1. Antibiotics, prepare concentrated stocks of (*See* **Note 3**):

a. 100 mg/mL ampicillin: dissolve 107 mg ampicillin sodium salt in 1 mL water. Filter sterilize using a PES syringe filter and store at –20°C.
b. 20 mg/mL tetracycline: dissolve 108 mg tetracycline hydroxide in 5 mL water. Filter sterilize using a PES syringe filter and aliquot stock in portions of 1 mL and store at – 20°C. Protect against light.
c. 50 mg/mL kanamycin: dissolve 120 mg kanamycin sulfate in 2 mL water, filter sterilize using a PES syringe filter, and store at –20°C.
d. 35 mg/mL chloramphenicol: dissolve 35 mg chloramphenicol in 1 mL 100% EtOH, filter sterilize using a PES syringe filter and store at –20°C.
e. 0.4 mg/mL trimethoprim: dissolve 40 mg trimethoprim in 100 mL water. Filter sterilize using a PES syringe filter and aliquot in portions of 1 mL and store at –20°C.
f. 1 mg/mL ofloxacin: dissolve 10 mg ofloxacin in 10 mL water. Filter sterilize using a PES syringe filter and aliquot in portions of 1 mL and store at –20°C.
g. 1.25 mg/mL rifampicin: dissolve 12.5 mg rifampicin in 10 mL water. Filter sterilize using a PES syringe filter and aliquot in portions of 1 mL and store at –20°C.
h. 10 mg/mL Carbonyl Cyanide m-Chlorophenylhydrazine (CCCP): dissolve 20 mg CCCP in 2 mL pure methanol, filter sterilize using a PES syringe filter and store at – 20°C.
2. LB medium: add tryptone (to a final concentration of 1% w/v), yeast extract (0.5% w/v) and sodium chloride (1% w/v) to a final volume of 500 mL using water. Autoclave for 15 minutes at 121°C and sterilize using a 0.22 μM PES filter to remove debris, which disturbs the flow cytometric analyses. Store at room temperature.

## 3. Methods

### 3.1. Generation of large fractions of tolerant cells

To investigate tolerant cells on the population level it is important to generate a cell population that consists of almost only dormant cells, despite a carbon source is available. In this section, we describe how to generate such a population, accomplished by a sudden nutrient shift. The procedure starts with generating a fully exponentially growing culture on glucose and continues with the steps to perform an abrupt switch to a different carbon source. In this section, fumarate is taken as the second carbon source because it generates the highest percentage of tolerant cells. Other gluconeogenic carbon sources can also be used for the switch (7). A schematic overview of this method is shown in Figure 1A, B and 1C.

1. Streak out an *E.coli* strain, e.g. K12, on LB-agar and incubate overnight at 37°C. Full-grown plates can be stored up to one month at 4°C sealed with parafilm.
2. Generate a culture that is fully exponentially growing on glucose, meaning that for several hours the cell number doubled at its maximal rate. Such a culture can be obtained by applying the following steps (*See* **Note 5**):

1. (Day 1) inoculate a 100 mL flask containing 10 mL M9 minimal medium supplemented with glucose to a concentration of 5 g/L with a single colony from an agar plate at around 5 p.m. Grow overnight at 37°C and shaking at 300 rpm till around 9 a.m. the next morning.
2. (Day 2) towards preparing the next preculture, first, determine the cell concentration in this overnight culture. For instance, dilute an aliquot of the culture 100x, transfer 200 μL to a 96-well plate (in case the used flow cytometer samples from 96-well plates) and measure the cell count (e.g. in 20 μL) with a flow cytometer (e.g. ACCURI C6, BD Biosciences) (*See* **Note 4**). Calculate the concentration in cells/mL, *C*, by using the formula:

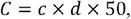

where *c* is the number of cells counted in 20 μL and *d* is the dilution factor. Then determine the volume of culture to be added to a fresh flask for having a target cell density of 3 · 10^8^ cells/mL at 5 p.m. on the same day as follows:

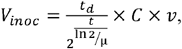

where *V_inoc_* is the volume that has to be added to the flask, *t_d_* is the target density (in cells/mL) at the end of the day, *μ* is the growth rate of the exponentially growing population (in h^−1^), *t* is time until the end of the day (in h) and *v* is culture volume (in mL). Add the calculated volume to a flask containing preheated (37°C) M9 minimal medium with 5 g/L glucose, and grow the culture at 37°C and shaking at 300 rpm.
3. (Day 2) towards performing the actual nutrient-shift in the next morning (day 3) with a culture that is fully exponentially growing on glucose, perform another subculturing step in the evening before (on day 2) to have 3·10^8^ cells/mL at the desired time point in the morning of day 3. To calculate the volume that has to be added to a new flask, repeat step 2b. Because of the huge dilution that has to be made for the inoculation of this overnight culture, it is recommended to inoculate several parallel flasks with dilutions such that one surely obtains at least one culture at the right cell density (*See* **Note 6**). Also, to prevent lag phase behavior, which can mess up the timing, it is recommended to use preheated and per-aerated media.
3. (Day 3) to start the nutrient shift, wait until the culture has reached OD_600_ of 0.3-0.8, first measure the cell concentration of the culture with the flow cytometer. The cell concentration should be around 2-5□10^8^ cells/mL. With the knowledge of the cell concentration, calculate the volume of the culture that contains 1.5□10^9^ cells. When staining is applied, skip the next step and go immediately to ‘Quantification of tolerant cells with membrane staining’.
4. Wash the cells to remove any residual glucose. Therefore:

1. Transfer the calculated volume to a 15 mL falcon tube.
2. Spin down the culture for 5 minutes at 3000g, 4°C, and discard the supernatant.
3. Wash the pellet with 5 mL ice-cold M9 minimal medium without carbon source by carefully pipetting. The ice-cold medium is used to slow down cells’ metabolism. Spin down, remove supernatant, and wash once more with the same amount of M9 minimal medium.
4. Spin down the culture for 5 minutes at 3000g, 4°C.
5. Discard the supernatant and resuspend the pellet in 1 mL M9 minimal medium without carbon source.
5. Transfer the cells to a 500 mL flask containing 50 mL preheated M9 minimal medium with 2 g/L fumarate. The cell concentration in the new flask is important because the initial cell density influences the fraction of cells that enter the tolerant state after the nutrient shift (*See* **Note 7**). With the *E. coli* wildtype strain BW25113, this procedure will result in about 99.99% of cells entering the tolerant state (7, 8). With other *E. coli* strains, this fraction can be different.

**Figure 1.**
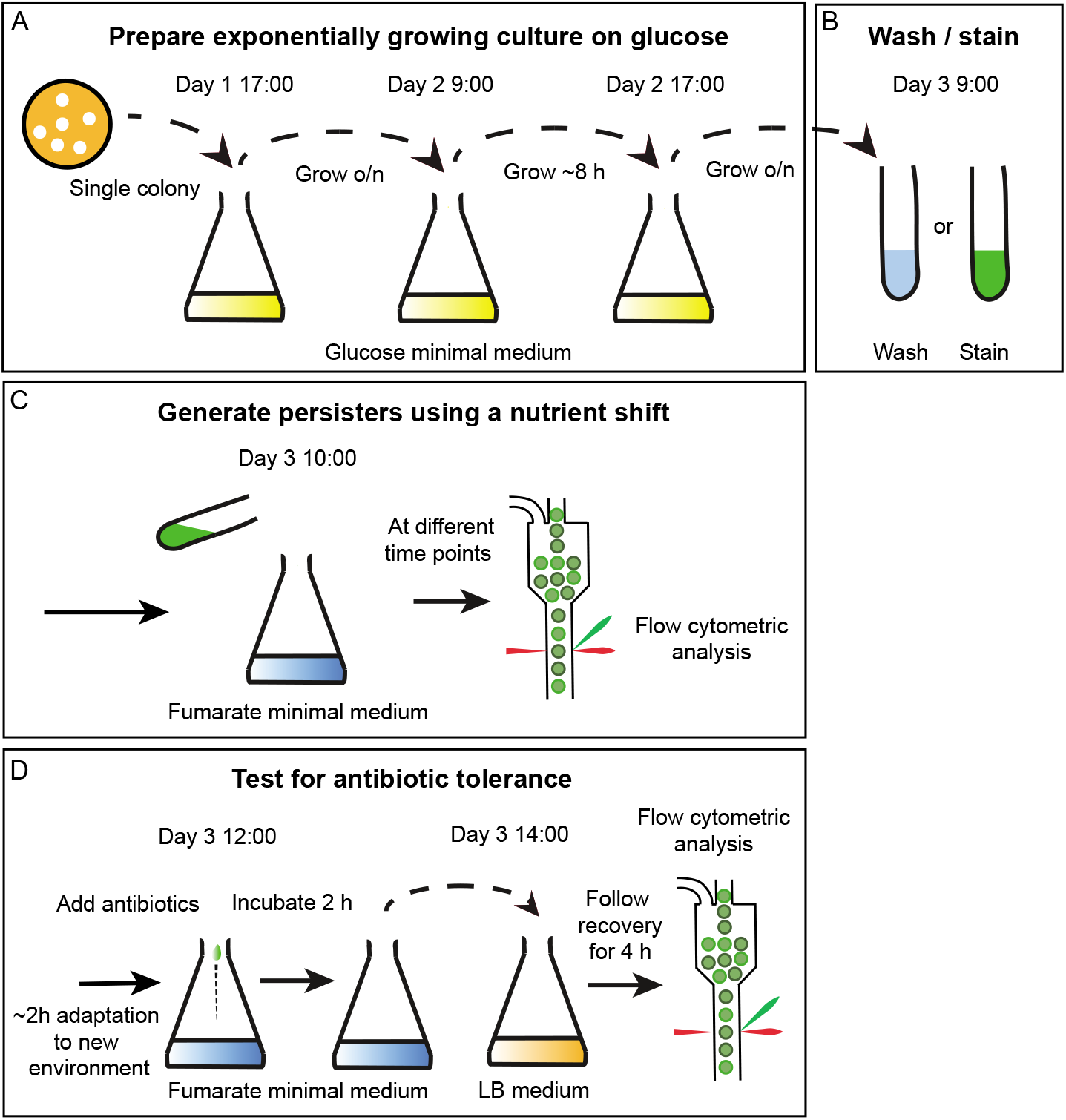
Schematic overview of the procedure for generating persisters and for testing for their antibiotic tolerance. (A) prepare an exponentially growing culture. Add a colony to 50 mL glucose minimal medium and grow overnight. Dilute this culture and grow during the day, dilute once more, and grow overnight. Make sure the culture is in an exponential phase when starting the staining. (B) wash or stain the cells to remove all residual glucose and optional, to stain the cells for tracking them with flow cytometry. (C) add the washed or stained cells to fumarate minimal medium and follow their growth at different time points using flow cytometric analysis. (D) to test antibiotic tolerance, add antibiotics after 2 hours in fumarate minimal medium. Incubate for 2 hours and transfer 500 μL to 50 mL LB medium. Follow regrowth for 4 hours using flow cytometry.

### 3.2. Identification and quantification of tolerant cells using membrane staining and flow cytometry

To determine the fraction of cells that enter the tolerant state, e.g. after the nutrient shift, one can stain the membrane of the cells with a fluorescent dye, can follow the fluorescence of the cell population after the nutrient shift over time and can analyze the resulting single-cell fluorescence data with a computational script. The dye we propose (PKH67, Sigma) stains the membranes of the cells and equally distributes over the two new cells after cell division. Thus, the fluorescence intensity of a cell halves with every division. Knowing the initial fluorescence intensity and comparing this with the fluorescence intensity at a later time point, one can estimate back how often a cell has divided (7). Specifically, to estimate the fraction of dormant cells, a Matlab script is used that fits a mathematical model to the time course data (i.e. cell counts, fluorescence intensity distributions). The model assumes that the fluorescence intensity of a cell decreases by half with each cell division and that cells have some autofluorescence. Further, it assumes exponential growth of the growing cells from some time point after the nutrient shift onward. As the fluorescence intensities of individual cells are not identical the model fits bimodal distributions to the fluorescence intensity distributions determined at the different time points and estimates the growth rates of the two populations of cells based on the total cell counts determined and fitted bimodal distributions. The model, its rational, and mathematics are explained in detail in our previous paper (7).

To determine the fraction of dormant cells that emerge after a sudden nutrient shift and to estimate the growth parameters (including among others, the growth rates of both populations), the membrane staining is applied in combination with the nutrient shift method to generate tolerant cells, meaning that the washing steps from the previous section are replaced by the following staining protocol. A schematic overview of this method is shown in Figure 1A, B, and 1C.

1. Generate exponentially growing cells on glucose according to steps 1 and 2 of the previous section.
2. Before the shift to fumarate, cells will be stained with a fluorescent dye. Before the staining, take Diluent C from the PKH67 kit from the fridge and let it warm up to room temperature. The dye should stay in the fridge.
3. Determine the volume of the glucose culture that contains 1.5□10^9^ cells. Transfer the calculated volume to a 15 mL falcon tube. Spin down the culture for 5 minutes at 3000g 4°C (Eppendorf centrifuge). Discard the supernatant very carefully by pipetting. Do not lose any cells, because the dye-to-cell number ratio is very important. Losing cells will increase the dye-to-cell ratio, through which cells could be stained too intensely, which in turn can lead to cell death. A sub-optimal dye-to-cell number ratio can also affect the viability of the cells resulting in fewer cells being able to recover after antibiotic treatment.
4. During the centrifugation as mentioned in the previous step, prepare a mix of 500 μL of room-temperature Diluent C with 10 μL of the dye solution (*See* **Note 8**).
5. **When performing steps 5 to 9 act fast and respect the timings** (*See* **Note 9**). Resuspend the cell pellet in 500 μL of solely Diluent C (room temperature) by carefully pipetting up and down. Make sure that the pellet is fully dissolved and that there are no droplets on the sides of the tube.
6. Add the prepared dye solution (10 μL of dye in 500 μL of Diluent C at room temperature) to the cell solution and mix by brief vortexing. Incubate the cells for exactly 3 minutes at room temperature (*See* **Note 10**).
7. Immediately after 3 minutes add 4 mL of ice-cold M9 medium with 1% (w/v) BSA and mix by brief vortexing (*See* **Note 11**).
8. Centrifuge the cells for 5 minutes at 3000 g and 4°C. Discard supernatant by pipetting, and resuspend the cells in 5 mL of ice-cold M9 without carbon source and centrifuge the cells again (5 minutes, 3400 rpm, 4°C).
9. Wash the cells once more in the same manner. Resuspend the cells in 1 mL room-temperature M9 medium without carbon source and transfer them to the preheated M9 minimal medium with 2 g/L fumarate (*See* **Note 12**) to yield an OD600 of approximately 0.6, which corresponds to a cell concentration of ~5 10^8^ cells/mL. Check the quality of the staining using flow cytometry by measuring a proper dilution of the culture. Well-executed staining should have only one peak on the FL-1 (533 nm) signal, and the cells should be 100-fold brighter than unstained cells.

### 3.3. Guide on how to use the Matlab script

The Matlab script and exemplary input files are provided on GitHub (https://github.com/molecular-systems-biology). The following steps describe the data acquisition and data format requirements.

1. To assess tolerant cells, it is important to generate tolerant cells as described in section 3.1: “Generation of large fractions of tolerant cells” and to stain them as described in section 3.2: “Identification and quantification of tolerant cells using membrane staining and flow cytometry”. The staining needs to be of an appropriate quality for the script to work optimally (*See* **Note 13**).
2. To obtain data that can be efficiently and easily used with the Matlab script, the measurements must be done at specific times. First, the first time point must be gathered as soon as possible after the switch.
3. The next data timepoint, which is the first data timepoint used to fit the model should be one in which the growing population is becoming visible in the data. This time point varies depending on the conditions and carbon sources used (*See* **Note 14**).
4. After taking these first time points, it is usually advised to make measurements every 30-60 minutes, depending on the growth rate of the growing population, until the growing population count is at least equal, or higher than the non-growing population count.
5. The data needs to be gathered over the growth of the culture (*See* **Note 15**). Certain factors need to be considered when gathering the data, for it to be usable with the script. The input data needs to be provided as a csv file containing the flow cytometry measurements for each cell at each time point, without a header (File 1). In the Matlab script, the following input needs to be provided:

a. The number identifying the column of the CSV file containing the relevant fluorescence intensity (FI) data (in case of our data, the number is 3 - variable SC).
b. The scaling factor. Different flow cytometers have different sensitivity and numerical output values. This factor is used to accommodate for these differences and make sure that the data from different machines can be used within the script capabilities. (variable maxval_new_FC).
c. The times when the samples were taken (variable tt)
d. The absolute cell concentration in the culture at the respective time points (variable cc)
e. The number of cells (or rows) in each CSV data file (variable g)
f. A specification of which of the data files should be used for the fitting. Depending on the nutrient switch, sometimes it takes hours before the growing population is numerous enough to be detected. The first timepoint used for the fit should have this population visible. Some data points can be excluded, for example, if the measurement has failed or has been inaccurate for any scientifically justified reason (variables indx and cc_indx).

In the Matlab script, one needs to define the allowed ranges of the parameters to be estimated as well as initial parameter guesses. These values should be close to the true values. Table 1 provides an overview of the parameters that need to be set and instructions on how to estimate these values, e.g. by visual inspection of the data or by using previous knowledge. It is advisable to run the script once on the data before setting the parameters. From the then generated “Fluorescence Data” figure (Figure 1), one can identify initial guesses (in matrix variable IG) and ranges for some of the parameters (in matrix variable bounds).

1. Two figures generated by the script are crucial at determining the quality of the fit, i.e. the figure “Cellcount curve fit check” and the multi-panel figure “Biggaussian fit for each time point”. In the Supplementary Figures, we show two examples of bad fits (files: badfit1, badfit2) and we show one example of a good fit (file: fit3). The files for these fits are provided with the Matlab script. The two figures show the cell count and fluorescence intensity data along with the plotted model predictions for both the growing and non-growing populations, and the sum of these populations. The way the model prediction fits the data can be used to assess whether the parameters are estimated correctly.
2. In the first bad fit (Supplementary Figure 1), the parameter bounds are set wrong. The FI means for both populations are overestimated, and the bounds are outside of the correct values. Moreover, the nongrowing population cell concentration is underestimated.
3. From the plots (Figure 2A), certain problems can be deduced:

a. The red line on the left panel that describes the modeled cell counts does not appear to fit with the cell count points from measurements. This is caused by the underestimation of the nongrowing population cell concentration.
b. The green line and the red line on the right panel do not fit the bimodal distribution. This is caused, on top of cause from problem 1, the overestimation of the FI means.
4. Moreover, we can see from the text output of the script that some of the parameters – the growth rate of the growing population in this case – reaches the lower limit of the bounds set, and thus it is not estimated correctly. However, we know what the growth rate of the used strain in given conditions should be and the fact that the estimate is equal to one of the boundaries is just a symptom of another issue, and not the issue itself.
5. If we fix the first problem and correct the nongrowing population cell concentration to a value that is close to the measured value, we obtain graphs shown in Supplementary Figure 2. While the graph on the left looks like it is a good fit, the graph on the right has the similar problems as before, stemming from the overestimation of the FI means. Fixing these values as described above, we can obtain a good fit.
6. It is clear from the plots shown in Figure 3B and 3C that the model closely fits the data and the data output can be trusted.
7. The parameters obtained from the fitting are output as text. The Matlab script used as an example generates the following data:

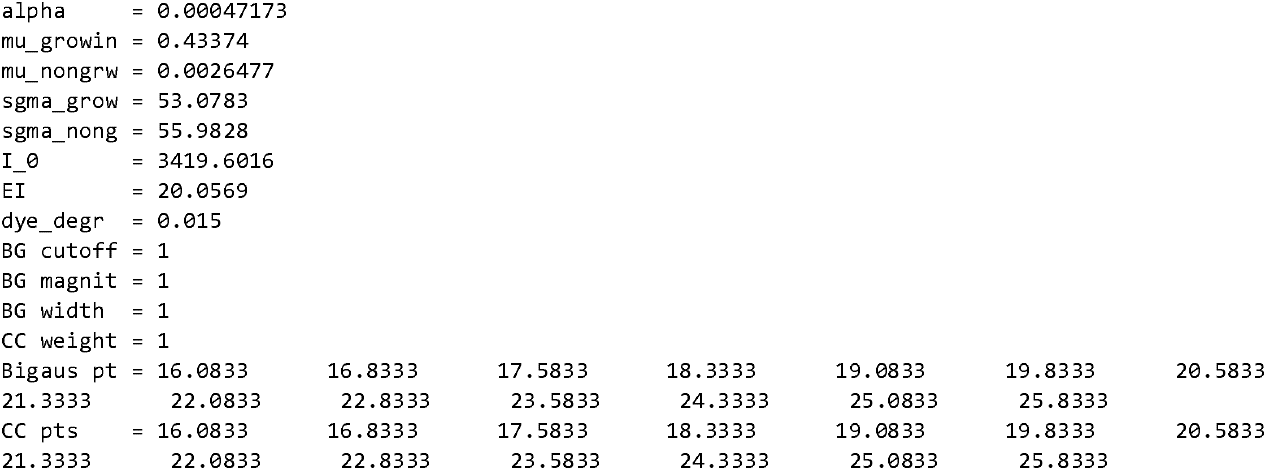
8. The most important obtained values are alpha (fraction of cells that entered dormancy after the nutrient shift), mu_growin (growth rate of the growing population), mu_nongrw (growth rate of the non-growing population). All these and other parameters, except alpha, should be checked against the lower and upper bounds set in the script and if they are equal to them, the bounds should be relaxed until they do not limit the results anymore.

**Table 1.** Overview of the parameters needed to be set for Matlab script. Left column: overview of parameters that need to be set. Right column: instructions on how to estimate these values.

| Parameter | How to determine |
| --- | --- |
| nongrowing population initial FI mean | From “Fluorescence Data” figure – see Figure 2A |
| nongrowing population FI stdev | From “Fluorescence Data” figure – see Figure 2A |
| nongrowing population cell concentration | The cell concentration after the nutrient switch is usually a good initial guess, except for nutrient switches in which the non-growing population is small. Upper and lower bounds can be very relaxed. |
| growth rate of non-growing population | From previous experiments |
| growing population initial FI mean | From “Fluorescence Data” figure – see Figure 2A |
| growing population FI stdev | From “Fluorescence Data” figure – see Figure 2A |
| initial growing population cell concentration | A guesstimate based on the expected number of adapting cells for a particular carbon source switch. Upper and lower bounds can be very relaxed. |
| growth rate of the population that starts to grow normally | From previous experiments determining growth rate on particular carbon source |
| unstained cell background autofluorescence | By analyzing data for unstained cells, or by checking the FI for very late samples, when cells have lost all their fluorescence. |

**Figure 2.**
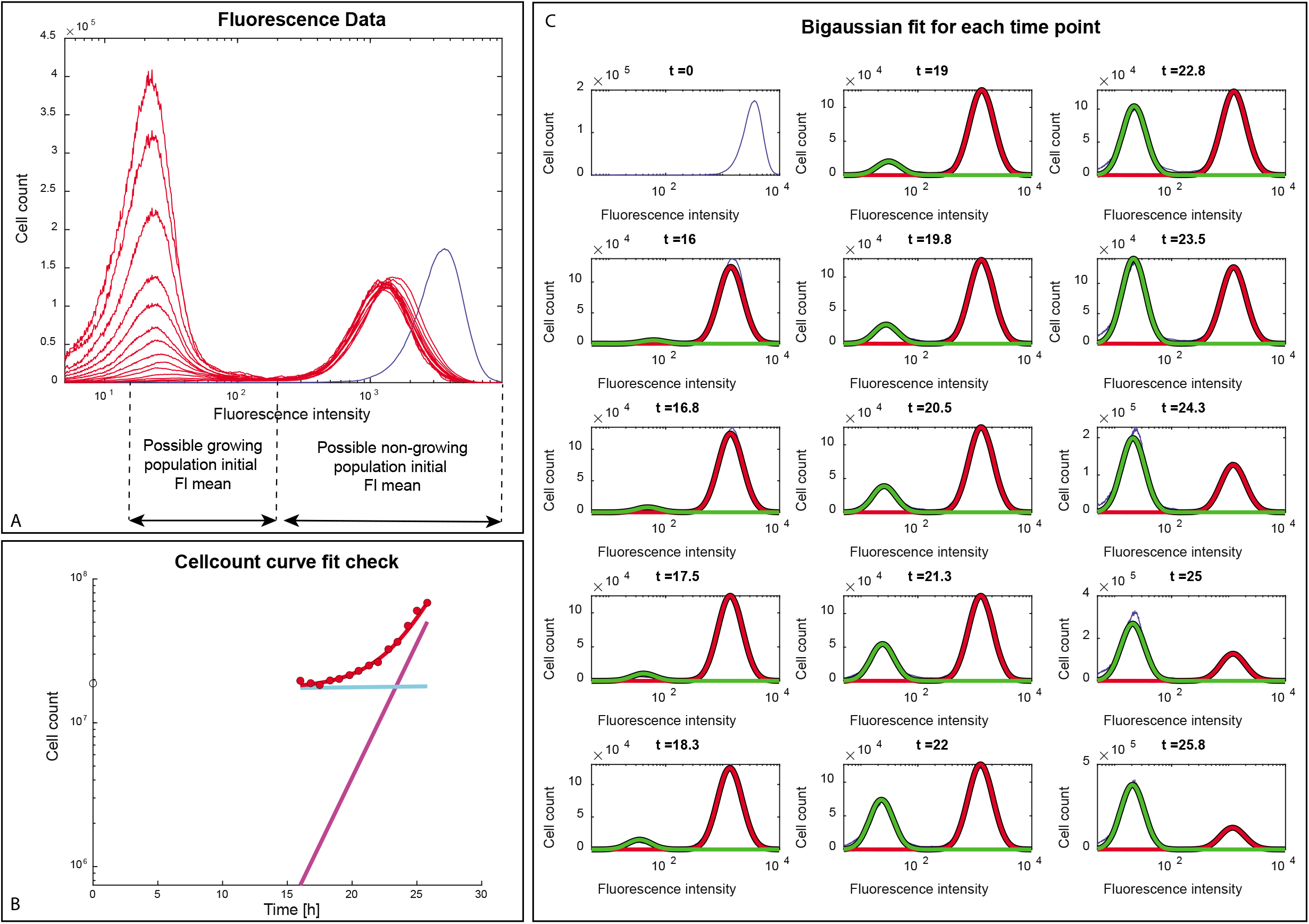
Finding initial parameters and example of a good fit. (A) How to find the initial parameter guesses using the Fluorescence Data figure generated by the script. The model needs a good estimation input to make it generating proper estimations. Therefore the mean fluorescence of the growing and the non-growing population needs to be estimated. To make a good estimation pick the average of the fluorescence in between the left arrows for growing cells and the average of the peak on the right for non-growing cells. (B) The cell count curve fit check. Empty disk – cell count at t = 0, red disks – cell counts used for the model fit, red line – predicted total cell count, cyan line – cell count of non-growing cells, magenta line – cell count of growing cells. (C) The model fit to the fluorescence data at each time point. Blue line – experimental data; green line – the distribution corresponding to the growing population; red line – the distribution corresponding to the non-growing population; black line – the sum of the distributions pictured by red and green line.

**Figure 3.**
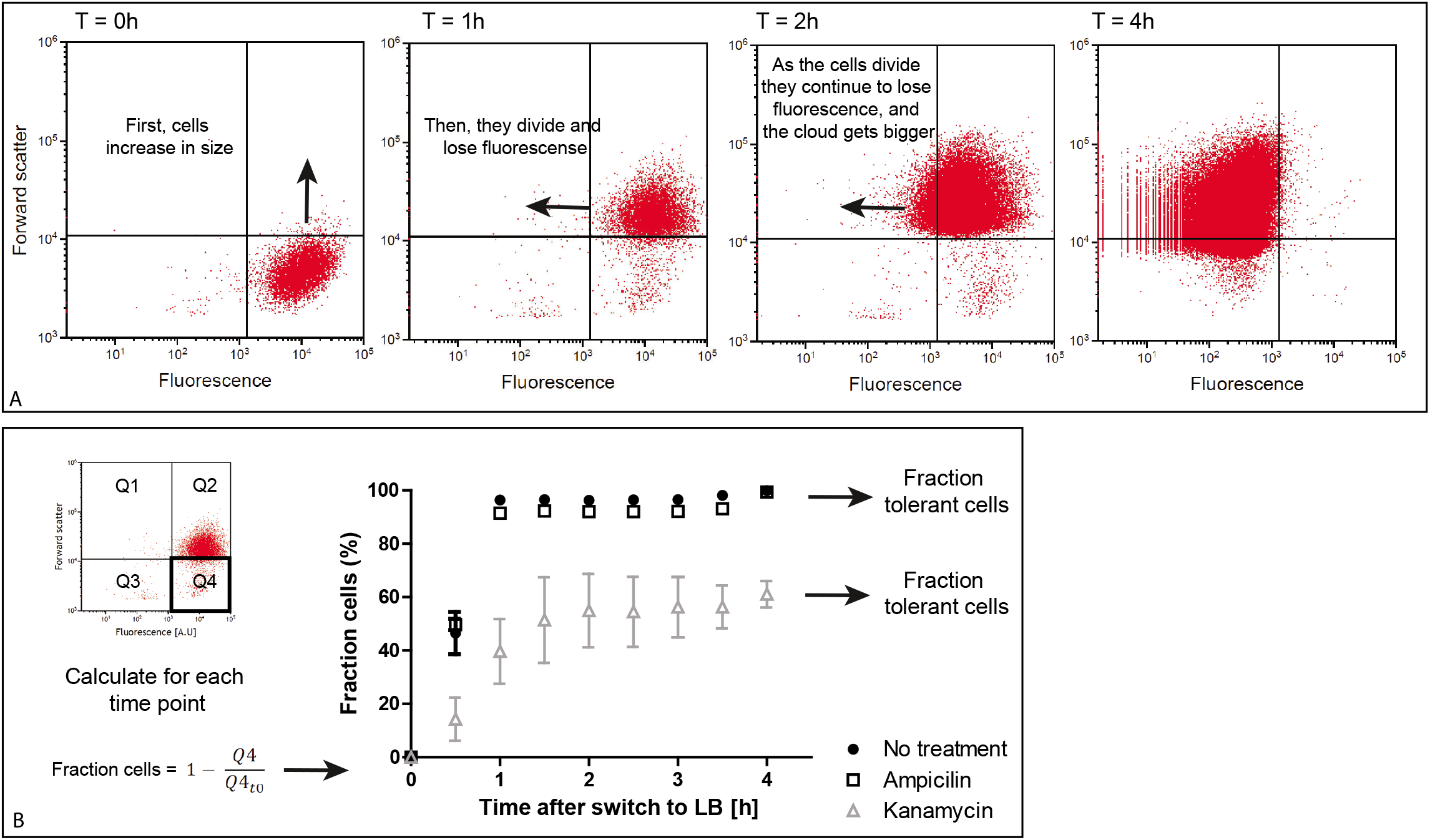
Example of regrowth in LB medium after antibiotic treatment. (A) Exemplary flow cytometry graphs over time: When tolerant cells (stained with fluorescent dye) are transferred to LB they first increase in size followed by a loss in fluorescence as a consequence of their divisions. Cells in Q1 are big and have lost their fluorescence. Cells in Q2 are big and have a high fluorescent intensity. Cells in Q3 are small and have lost their fluorescence. Cells in Q4 are small and are fluorescent. (B) Left, the formula of how the fraction of tolerant cells is calculated. Right, an example graph of treatment with 2 different antibiotics. The fraction of cells for each time point is calculated by 1 minus the number of cells in Q4 divided of the number of cells in Q4 on timepoint 0. Cells that are killed by antibiotics will not regrow in LB medium and therefore will not leave section Q4 in the flow cytometer graph. After 4 hours a steady state is reached and the fraction of cells on T = 4 can, therefore, be used as the ultimate fraction of viable cells.

### 3.4. Assessment of antibiotic tolerance with flow cytometry

The ability to regrow after antibiotic treatment is an indication that a cell has been in the tolerant state during treatment. While typically, such regrow experiments are done with plating assays (i.e. determination of colony-forming units, CFUs), here, we proposed a method to perform such regrow assessments with flow cytometry. These assays require the above-mentioned membrane staining. Despite the staining procedure that needs to be carried out, this technique is less laborious and has less variability than plating assays (9). Also, it was found that more cells were able to wake up after dormancy when liquid media is used compared to regrowth on plates (10). A schematic overview of this method is shown in Figure 1D.

1. To assess the antibiotic tolerance of cells, it is important to generate tolerant cells as described in section 3.1: “Generation of large fractions of tolerant cells” and to stain them as described in section 3.2: “Identification and quantification of tolerant cells using membrane staining and flow cytometry”.
2. After the transfer to the fumarate medium, it takes a certain while until the non-adapting cells to develop full tolerance against antibiotics. It is recommended to let the cells adapt for at least 2 hours after staining (*See* **Note 16**) before treating them with antibiotics.
3. After >2 hours, add the antibiotics. We tested ampicillin (100 μg/ml), tetracycline (20 μg/mL), kanamycin (100 μg/ml), chloramphenicol (140 μg/ml), trimethoprim (5 μg/mL), rifampicin (100 μg/ml), ofloxacin (5 μg/ml) and CCCP (50 μg/ml) (8). Concentrations were obtained from literature and from survival assays on cells growing on glucose (*See* **Note 17**). Because some cells undergo a reductive division after the switch to fumarate, the number of cells after 2 hours is slightly higher than at time point zero. Therefore, it is required to measure a dilution of the culture direct after staining and right before antibiotics are added.
4. Incubate the cultures that now contain the antibiotics for 2 hours whilst shaking at 300 rpm at 37°C. After the incubation time, measure again a dilution of the culture with the flow cytometer to determine the cell count. The cell count should not have increased after the addition of antibiotics. In some cases, the cell count can even be decreased, e.g. when a bacteriolytic antibiotic is used (*See* **Note 18**).
5. To assess the fraction of cells being able to recover antibiotic treatment the antibiotics are diluted out. Pipette 500 μL of the culture into 50 mL preheated and filtered LB medium, mix well and transfer a sample of the non-diluted culture in the 96 wells plate (from where the flow cytometer will take the sample) and measure it straight away with the flow cytometer. The sample should not be diluted because there are now 100-fold less cells in the culture than in the flask where the cells were incubated with the antibiotics. LB contains, when only autoclaved, a lot of components that lead to a strong background signal in the flow cytometer. This will interfere with your sample because the debris will appear around the same size as your cells on the FSC-SSC dot plot and therefore it is required to use filtered LB medium to lower the background signal. The first sample taken from the LB culture is very important because it reflects all cells in their dormant state and it is used to calculate the fraction of cells that became dormant after the nutrient-switch (*See* **Note 19**).
6. From now on, subject the LB cultures to flow cytometric analyses every 30 minutes for a period of 4 hours. Cells have a short doubling time on LB and thus cell counts can quickly enter a range that is beyond the linear range of the flow cytometer. Take notice of the cell count in each sample and determine the appropriate dilution for the next sample (*See* **Note 20**).
7. The fraction of tolerant cells is calculated by subtracting the number of nondividing cells (cells that do not exit the bottom right part of Figure 3A) in each time point from the initial number of nondividing cells.
8. Samples measured directly after staining, after the addition of antibiotics, and right before the nutrient shift are solely used as quality controls for the experiment. The cells should show one single small peak directly after the switch to fumarate and the population should have significantly slowed down growth before antibiotics have been added. After the two hour treatment with antibiotics the growth should have completely stagnated or the cell number could even have dropped in case a bacteriolytic (e.g. ampicillin) has been used.

#### 3.4.1 Data analysis

1. For data analysis of the total fraction of tolerant cells, the cells are followed for four hours after the switch to LB, and the loss of fluorescence as well as the size of the cells is used to determine the fraction of the population being able to escape dormancy after antibiotic treatment. It is required to set up a dot plot to gate the cells (FSC-H vs SSC-H) and a dot plot using cell size and fluorescence (FL1-A vs FSC-A).
2. To determine the fraction of cells escaping dormancy the number of cells transferred to LB is taken at time point zero (T0). For each following time point, the number of nondividing cells is subtracted from the cells transferred at T0. It is not possible to use the number of escaping cells since they start dividing after the switch to LB (*See* **Note 21**) as shown in Figure 3A.
3. To define the non-growing population, the quadrant function. As in Figure 3A, it needs to be adjusted manually for each sample. Particularly in samples where a big part of the population just started to divide it can be hard to distinguish the non-growing population from the growing population. It is advised to use the later time-points to set the quadrant and use the same settings for the early division samples (*See* **Note 22**). When one is only interested in the final number of tolerant cells and not in the recovery dynamics one can decide to not analyze these samples. An example of the recovery dynamics is shown in Figure 3B.
4. The data from 4 hours on LB is used to calculate the final fraction of tolerant cells. Take the number of nondividing cells and subtract it from the number of cells added to LB at T=0. It is optional to take the average of the last two, or the last three measurements to calculate the final fraction of tolerant cells (only if the recovered fraction reached steady state). Averages and standard deviations are calculated from three individual experiments and shown as box plots.

## 4. Notes

1. If antibiotics are required for the particular *E.coli* strain it is important that the antibiotic is always added after autoclaving the LB-agar and after the LB agar has cooled down to ~55°C, to avoid degradation of the antibiotic. It is recommended to prepare 1000x concentrated stocks of the required antibiotics, to store them at –20°C and thaw them just before pouring plates.
2. When using the glucose-to-fumarate rapid nutrient shift to generate large fractions of tolerant cells, it is important to not use cellulose acetate filters because these filters could release acetate into the medium (13), which *E. coli* can use as a carbon source. As *E. coli* can have different preferences on which gluconeogenic carbon source to use first, even a small concentration of acetate could affect the results. PES filters are equally priced and do not release any compound that can act as a substrate for *E. coli*.
3. The antibiotic concentrations used in the working solutions are based on literature research and on experiments where we tested the antibiotics on *E.coli* cell growing on fumarate. The concentration of the antibiotic stocks is based on the solubility of the antibiotic. For ampicillin and tetracycline, the solubility is very high so it was decided to make a 1000x concentrated stock.
4. When measuring and counting *E. coli* cells, the flow cytometer needs to be calibrated for small cells. Typically, the default settings are not suitable for *E. coli* cells. With the BD Accuri C6, the threshold for *E. coli* should be 8000 for the forward scatter (FSC-H) and 500 for the side scatter (SSC-H). Further, debris in the medium can disturb the measurement. Thus, it is required to only use filtered media when measuring samples with the flow cytometer. For reliable cell counts with the Accuri C6, the events measured by the flow cytometer should be between 10 000 and 100 000 cells in 20 μL of the measured sample. This level ensures a high enough amount of cells in comparison to machine noise, and low enough to not cause over-saturation of the instrument.
5. For the successful generation of almost 100% tolerant cells with a glucose-to-fumarate shift, the cells in the glucose culture must be in a fully exponentially growing state on glucose for at least 24 hours before the nutrient shift. Because cells typically have a lag phase after inoculation from the LB plate, we recommend starting culturing cells at least one day before the actual nutrient shift to ensure the maximal growth rate on glucose.
6. To ensure cells that cells keep on growing exponentially also after the re-inoculation, it is crucial to pre-heat the media before dilution of the cultures. But even then, typically short lag phases occur after diluting, and it is thus recommended to not only inoculate the calculated number of cells but also to prepare a preculture starting with 2x the number of cells required, such that at the desired time point in the morning one surely obtains a culture with the proper cell density.
7. Previous research has shown that the initial cell density after the nutrient shift influences the fraction of dormant cells after nutrient shift (14). The absolute number of adapting cells seems to be constant regardless of the initial cell density after a glucose to fumarate shift, resulting in a higher fraction of cells adapting when a flask is inoculated with a low cell density (13). Furthermore, note that with different *E. coli* wildtype strain strains different fractions of dormant cells can emerge.
8. Preparing this mixture too soon will cause the dye to clump and staining intensity will be sub-optimal. The solution can be prepared during the first centrifugation step, although the Diluent C must be brought to room temperature earlier. When multiple samples are stained in parallel, always change the pipette tip to prevent water transfer to the vial containing the dye stock. The dye-to-cell number ratio is very important for proper staining. The ratio is good, when the cells clump a bit during staining, as indicated by a slightly cloudy solution.
9. Leaving droplets on the side on the tube will create a fraction of unstained cells because droplets containing cells do not get in touch with the dye. This will be seen as a separate unstained fraction and will influence your results or even make them unusable.
10. Longer incubation will cause the cells to die, whereas shorter incubation will cause the cells to be stained sub-optimally. If you stain cells of multiple different samples in parallel, add the dye in 20-30 second intervals, such that the incubation time can be strictly adhered to.
11. BSA blocks the remaining dye molecules, thereby preventing the cells from being killed. It is recommended to use a timer and make sure you have the M9 medium with BSA in your pipette ready so that you only have to release it from the pipette into the tube containing the cells when the 3 minutes incubation time is over.
12. It is recommended to prepare the flasks with preheated and pre-aerated M9 medium with 2 g/L fumarate before harvesting the cells.
13. To obtain a good fit of the data to the model and thus reliable parameter estimates, the data needs to fulfill certain criteria. First, the fluorescence intensity of stained cells should be at least two orders of magnitude higher than the background fluorescence of unstained cells. This, in turn, means that the fluorescence intensity decrease can be tracked over 5-6 divisions, which is enough for most carbon source switches. Moreover, the fluorescence intensity data of stained cells should form a single peak in the histogram. The narrower the peak, the better, as the growing and non-growing populations will be better separated from each other in the obtained data. While in our experiments we had the best results using the green fluorescent dye, we have also used a red fluorescent dye (PKH26), which was giving us data of lesser quality. In principle, any product that utilizes a fluorophore linked to an aliphatic chain that intercalates in the cell membrane could be utilized to generate the data for the script, and there are many alternative products available on the market that have different excitation and emission properties that might be best suitable for the equipment used. However, due to different protocols and properties of these dyes, appropriate controls would have to be made to exclude the effect of the staining procedure on the obtained results.
14. The higher the fraction of adapting cells, the sooner the first usable data point can be taken. In the case of a glucose-to-fumarate switch, 15-16 hours after the switch is usually a good starting point, as the growing population starts to be large enough to be detectable, while it still has a fluorescence intensity that is above the autofluorescence. Capturing time points, in which the growing population still has some fluorescence above the cellular autofluorescence is crucial for the script to estimate the fraction of growing cells accurately.
15. The fluorescence intensity data from the flow cytometer needs to be in log10 space. Depending on the cytometer used, a transformation needs to be done on the data (as we do in case of the Accuri C6 cytometer) or is already done in the cytometer or its software. Moreover, depending on the flow cytometer, the numerical range of values can be different and this needs to be addressed by setting the scaling factor.
16. After transferred to the fumarate medium, several cells will undergo a reductive cell division within about the first hour. Experiments have shown that after 2 hours on fumarate, all cells of the population become tolerant against ampicillin (8). Adding antibiotics too early might result in fewer cells surviving treatment, whereas incubating longer than 2 hours is not increasing the tolerant population.
17. For each antibiotic used in our experiments, we checked in the literature which concentrations were used in *E.coli* inhibition experiments. We used this information as a starting point to explore which concentration of each antibiotic kills growing cells. Before the tolerance experiments were done we carried out identical experiments with glucose grown cells to determine the concentration of antibiotics needed to kill a growing population.
18. When switching the cells to fumarate 0.01% of the population will adapt and start proliferating (7). Those cells will be sensitive to antibiotic treatment. Especially when the tolerant cells are kept for longer periods (~24 h after the nutrient shift) a significant population of growing cells is visible which are sensitive to antibiotics. When a bacteriolytic antibiotic such as ampicillin is used, the treatment will cause these cells to lyse. Therefore, the cell count after treatment can be lower than before treatment.
19. To check cells’ ability to survive antibiotic treatment they are transferred to LB medium. No washing steps are applied to prevent the cells from extra stress. By only transferring 500 μL in 50 mL the antibiotics are diluted 100 fold, enough to nullify their inhibiting effect.
20. Bacteria grow fast on LB medium, μ = 1.9 h^−1^ (15). Therefore, it is essential to take regular measurements, to generate reliable data points. Dead cells do not proliferate, meaning that their number will remain the same and they will not lose their fluorescence. They will be visible as a stained cloud of small cells in the FL1-A versus forward scatter (FSC-A) plot. However, when the descendants of the small growing population keep increasing their number, they will outgrow the linear range of the flow cytometer and a bigger dilution must be made to measure the sample. This has the risk to lose the visibility of the population of dead cells because they excessive diluted and their cell count is not reliable. In particular, when only a small fraction of cells are not dividing a small dilution will introduce a big measurement error by dilution out the number of non-growing cells.
21. Bacteria escaping the dormant state do this by resuming their growth. Since it is impossible to determine the individual growth rate for each cell there is no way to use the growing population for calculating the fraction of cells able to escape dormancy. However, since we know the original number of cells transferred to LB and we can distinguish growing from non-dividing cells we can use the number of non-dividing cells and the number of cells at T0 to calculate the fraction of the population which has survived antibiotic treatment.
22. When cells start waking up from their dormant state, they will first get bigger, followed by the loss of fluorescence. This can be seen in a dot plot as in Figure 3A (FL1-A vs FSC-A). However, in some cases, the cells may wake up a bit slow, and in the earlier time-points of the recovery experiment, the population of cells increasing size might overlap the nondividing population. In that case, it is advised to determine the non-dividing population at a later time-point and use those settings in the more indefinite sample.

## Acknowledgment

This work was supported by the Netherlands Organisation for Scientific Research (NWO) through a VIDI grant to MH [project number 864.11.001].

---

**Supplementary figure 1 – Bad fit 1**. (A) The cell count curve fit check. Empty disk – cell count at t = 0, red disks – cell counts used for the model fit, red line – predicted total cell count, cyan line – nongrowing cell count, magenta line – growing cell count. (B) The model fit to the fluorescence data at each time point. Blue line – experimental data; green line – the distribution corresponding to the growing population; red line – the distribution corresponding to the non-growing population; black line – the sum of the distributions pictured by red and green line.

**Supplementary figure 2 – Bad fit 2.** (A)The cell count curve fit check. Empty disk – cell count at t = 0, red disks – cell counts used for the model fit, red line – predicted total cell count, cyan line – nongrowing cell count, magenta line – growing cell count. (B)The model fit to the fluorescence data at each time point. Blue line – experimental data; green line – the distribution corresponding to the growing population; red line – the distribution corresponding to the non-growing population; black line – the sum of the distributions pictured by red and green line.

## Notes

### Competing Interest Statement

The authors have declared no competing interest.

