## Supplementary figures and images for "A robust method for generating, quantifying and testing large amounts of *Escherichia coli* persisters"

### Supplementary Figure 1

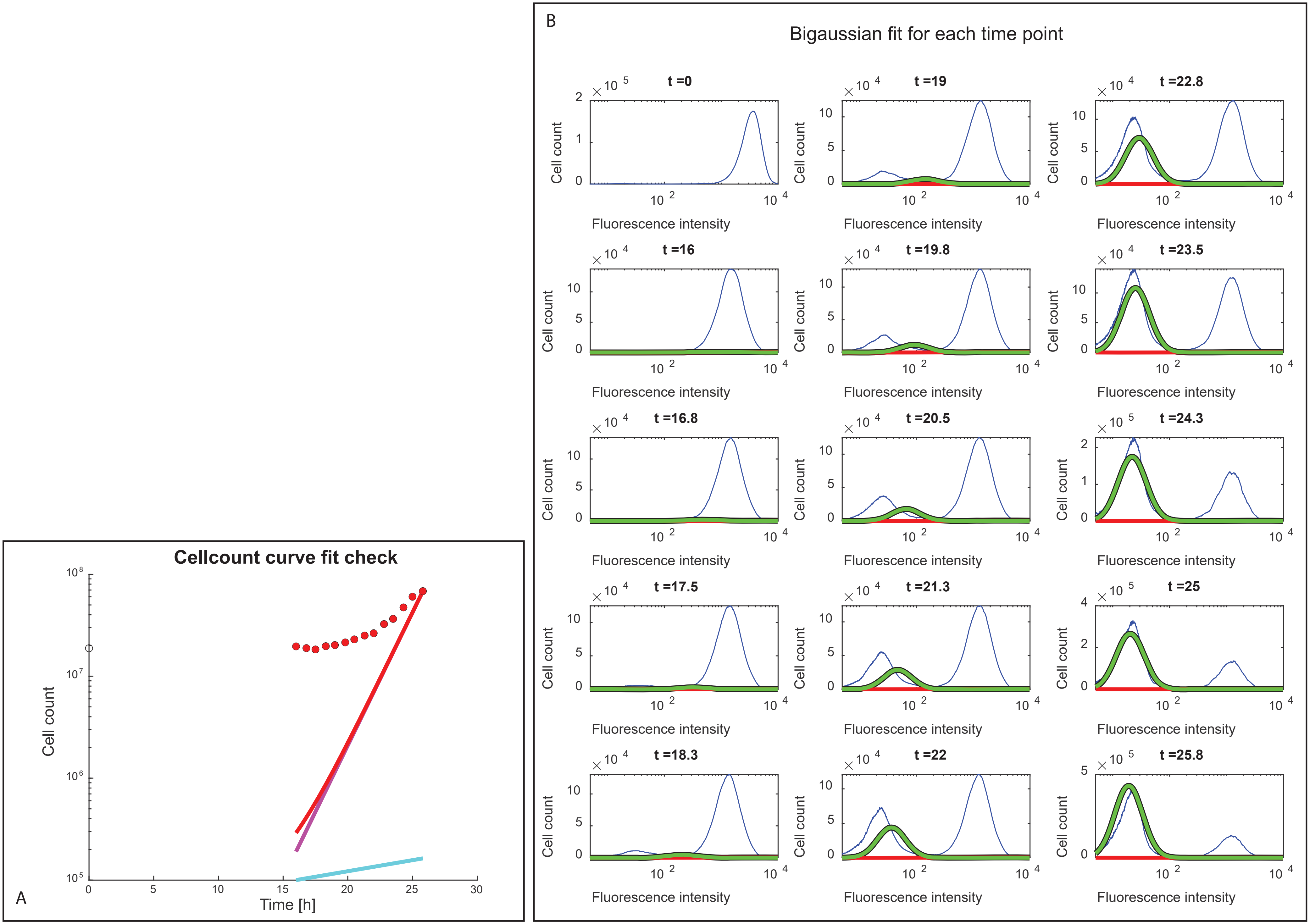

### Supplementary Figure 2

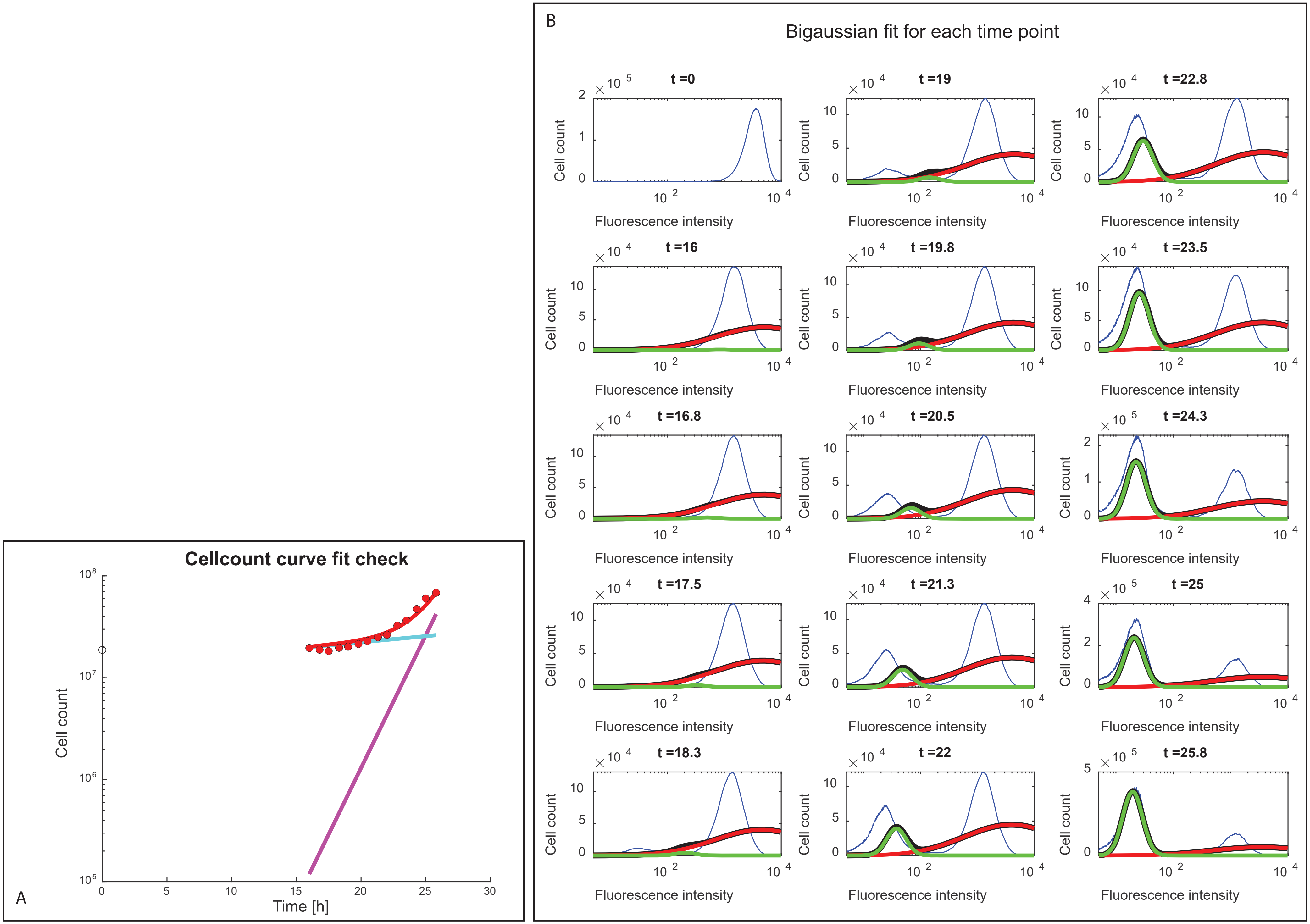
